# Evolution of highly repetitive silk genes in the Luna moth, *Actias luna*

**DOI:** 10.1101/2025.08.07.669095

**Authors:** Bert Foquet, Lauren E. Eccles, Amanda Markee, Deborah A. Triant, Paul B. Frandsen, Whitney L. Stoppel, Akito Y. Kawahara

**Affiliations:** McGuire Center for Lepidoptera and Biodiversity, Florida Museum of Natural History, University of Florida, Gainesville, FL 32611, USA; Department of Chemical Engineering, University of Florida, Gainesville, FL 32611, USA; American Museum of Natural History, New York, NY 10024, USA; Data Analytics Center, Research Computing, University of Virginia, Charlottesville, VA 22903, USA; Department of Plant and Wildlife Sciences and Bean Life Science Museum, Brigham Young University, Provo, UT 84602, USA

**Author notes:** Author for Correspondence: Bert Foquet*, ^1^McGuire Center for Lepidoptera and Biodiversity, Florida Museum of Natural History, University of Florida, Gainesville, FL 32611, USA.

**Keywords:** Gene evolution, Lepidoptera, repetitive DNA, sericin, Saturniidae

## Abstract

Gene duplications are a major driver of molecular diversification and phenotypic evolution. Arthropod silk genes provide an excellent model for studying these processes due to their highly repetitive sequences and rapid evolutionary rates. In Lepidoptera, the *Fibroin heavy chain* (*fibH*) gene encodes the primary structural protein for silk fibers, contributing largely to their mechanical strength. This inner fibroin core is surrounded by an outer coating composed primarily of sericins. Sericins are a group of highly repetitive, serine-rich proteins that modulate silk fiber properties. Although sericins in the Domesticated silkworm (*Bombyx mori*) have been associated with life stage-specific variation in silk characteristics, their evolution and function across Lepidoptera remain poorly understood. Here, we provide a detailed molecular characterization of sericin genes in the Luna moth (*Actias luna*), a saturniid species known for forming dense, robust, silk-woven cocoons. We identified eight sericin genes that (1) are frequently arranged into clusters of closely related paralogs, (2) exhibit considerable variation in repeat number and amino acid composition, and (3) display distinct gene expression patterns across life stages. A comparison of sericin genes across Saturniidae and Bombycidae reveals evidence for convergent subfunctionalization. These findings suggest that sericin gene duplications enable dynamic shifts in silk composition both within and between species, potentially reflecting adaptive responses to ecological and functional demands.

**Significance Statement:** Gene duplications are thought to be a major driver of molecular diversification and phenotypic evolution. Arthropod silk genes, characterized by their repetitive sequences and rapid evolution, provide an ideal model for studying these processes. Sericins, a group of highly repetitive, serine-rich silk proteins, are hypothesized to have contributed to the diversification of silk properties, both within and across lepidopteran species. However, their diversity and evolution is poorly understood. Focusing on the Luna moth (*Actias luna*), we show that sericin gene duplications across Saturniidae have led to subfunctionalization, enabling changes to silk composition. These modifications may represent adaptative responses to ecological and functional demands.

## Introduction

Gene duplications are one of the primary drivers of phenotypic diversity, producing redundant gene copies that can evolve without compromising their original role (Birchler & Yang 2022; Ding et al. 2012; Kuzmin et al. 2022). Although duplicated genes are often lost, others are retained and may acquire novel functions (neofunctionalization), partition ancestral roles (subfunctionalization), increase gene dosage, or act as compensatory backups (Birchler & Yang 2022; Kuzmin et al. 2022). These mechanisms have contributed significantly to shape major evolutionary innovations, from the diversification of animal body plans (Wagner et al. 2003) and variation in human traits (Conrad & Antonarakis 2007), to the emergence of silk fiber diversity in spiders (Garb et al. 2010). While gene duplications can arise from non-homologous recombination repair (Koszul et al. 2004) and transposition events (Cerbin & Jiang 2018; Hughes et al. 2003), most occur in regions rich in repetitive DNA (Babcock et al. 2003; Delihas 2020; Dennis et al. 2017). In these regions, non-allelic homologous recombination (Taylor et al., 1957) and unequal crossing over during meiosis (Smithies 1964) are the predominant drivers. These processes can also act within individual repeat-rich genes, accelerating their evolution and leading to the development of novel traits (Cheng & Chen 1999; Delihas 2011; King 2024). Despite their evolutionary importance, such genes have been historically difficult to study due to their complex structure. However, advances in long-read sequencing technologies now make it possible to resolve these regions with high accuracy at the genomic level (Hotaling et al. 2021; Kawahara et al. 2022; Krsticevic et al. 2015).

Arthropod silk genes, known for their extreme repeat content, have emerged as a powerful model to investigate the evolutionary dynamics of repetitive, protein-coding DNA. These genes encode for silk fibers that have captivated human societies for millennia due to their remarkable mechanical properties, combining strength and extensibility, which arise from their semicrystalline structures and long, repetitive protein sequences (Aikman et al. 2025; Sehnal & Sutherland 2008). The major silk proteins, including fibroins in butterflies, moths, and caddisflies (Amphiesmenoptera) and spidroins in spiders (Araneae), have convergently evolved highly repetitive sequences rich in alanine, glycine, and serine (Gatesy et al. 2001; McKim & Turner 2024; Sutherland et al. 2010; Walker et al. 2012). The spidroin gene family has undergone extensive gene duplication, shifts in expression, and diversification of its repetitive domains, collectively enabling the unparalleled mechanical diversity of spider silk (Clarke et al. 2017; Kono et al. 2019; Starrett et al. 2012). In contrast, duplications of the *Fibroin heavy chain* (*fibH*) gene – the primary silk gene in Lepidoptera and Trichoptera (Heckenhauer et al. 2023; Yonemura et al. 2009; Zhang et al. 2024) – are rare. This suggests that in Lepidoptera, other genetic mechanisms might contribute to previously reported functional and structural silk diversity (Eccles et al. *Under Review*; Guo et al. 2022; Peng et al. 2019).

Lepidopteran silk fibers consist of an inner core, consisting primarily of the fibH protein, and an outer coating. This outer coating makes up 20–50% of the silk fiber and functions as an adhesive and protective layer influencing fiber structure, stability, adhesion and mechanics (Dong et al. 2013; Gheysens et al. 2011; Guo et al. 2022; Malay et al. 2016; Peng et al. 2019; Takasu et al. 2017; Wu et al. 2022). Sericins, a group of distantly related, serine-rich and highly repetitive proteins, make up the largest portion of this outer coating (Dong et al. 2013; Guo et al. 2025, 2022; Rouhová et al. 2024). In the domestic silkworm, *Bombyx mori*, only three of six known sericins are present in cocoon silk, while the other three are found in silks produced during early larval stages (Dong et al. 2013, 2019; Guo et al. 2022; Peng et al. 2019; Wu et al. 2024). These stage-specific expression patterns are correlated with variation in silk mechanics and properties across developmental stages (Guo et al. 2022; Kludkiewicz et al. 2009; Takasu et al. 2010), suggesting that the sericin-rich outer coating, rather than the fibroin core, determines the functional diversity of silk. Comparative studies of sericins, both between and within species, are complicated by their high sequence divergence (Kmet et al. 2023; Tsubota et al. 2021; Wu et al. 2022), leaving the evolutionary history of this gene family poorly resolved. As a result, it remains unclear whether other lepidopteran species share the sericin expression patterns observed in *B. mori*.

Moths in the family Saturniidae diverged from the lineage leading to *B. mori* approximately 70 million years ago (Kawahara et al. 2019), making them well-suited for investigating silk gene diversification in a non-model system. Saturniids have long attracted interest as alternatives to *B. mori* due to their large body size and robust cocoons, which are the source of several commercially and culturally important wild silks, including tussah, muga, and eri. While the silk properties of these species are relatively well characterized (Holland et al. 2012; Malay et al. 2016; Reddy & Yang 2012; Schmidt et al. 2023), and silk genes have been identified in several Saturniidae (Rouhová et al. 2024; Tsubota et al. 2016; Žurovec et al. 2016), little is known about the genomic organization, variability, and evolutionary history of sericins.

The Luna moth (*Actias luna*), a large and iconic North American saturniid, produces silk throughout multiple larval stages, including anchoring silk, molting pads, and a strong silk cocoon for protection (Eccles et al. *Under Review*; Reddy & Yang 2012; Chen et al. 2012a). A recently published, high-quality genome assembly (Markee et al. 2024), along with a detailed analysis of its silk structure (Eccles et al. *Under Review*), makes *A. luna* an ideal species to characterize the structure, expression, and evolution of sericin genes in a non-model lineage. In this study, we identified and extracted eight sericin genes in *A. luna*, examined their genomic locations, and encoded protein sequences to sericins from closely related saturniids, including *Actias selene*, *Antheraea assamensis*, *A. pernyi*, *A. yamamai*, *Hyalophora cecropia*, *Rhodinia newara*, *Samia ricini*, as well as *B. mori*. We also examined changes in sericin gene expression across developmental stages and assessed variation in predicted protein composition among the *A. luna* sericins.

## Results

### Identification of sericins in the *Actias luna* genome

We applied three different methods to identify sericin genes in the *A. luna* genome: sequence similarity with known sericins, shared sequence motifs with known sericins, and genetic proximity to other *A. luna* sericins. First, we generated a sericin dataset containing 22 previously characterized sericins from seven species of Saturniidae (*Actias selene*, *Antheraea assamensis*, *Antheraea pernyi*, *Antheraea yamamai*, *Hyalophora cecropia*, *Rhodinia newara*, and *Samia ricini*) and *B. mori*, a member of the closely related Bombycidae (Dong et al. 2015, 2019; Guo et al. 2022; Rouhová et al. 2024; Takasu et al. 2007; Tsubota et al. 2016; Žurovec et al. 2016), **Supplementary Table S1**). This dataset was used to query the *A. luna* genome for sericins. Second, a set of conserved sequence motifs was generated by aligning all sequences in the sericin gene dataset and was used to further unveil luna moth sericin genes. Both methods yielded the same set of six sericins (**Table 1**). Two more putative sericins were identified by a targeted search for sericin-like genes in genomic proximity to these six sericins (**Table 1**). To avoid any assumption of homology with *B. mori sericin 1-5*, these eight *A. luna* sericins were given letter codes based on their order in the genome (*sericin A-G*; *serA-G*), except for *A. luna sericin 1* (*ser1*) which we found to be a homologue of *B. mori sericin 1* (**Table 1**). The eight *A. luna* sericin genes share a set of structural characteristics previously described from sericins (Dong et al. 2019; Garel et al. 1997; Guo et al. 2022; Kludkiewicz et al. 2009; Takasu et al. 2007; Wu et al. 2024; Žurovec et al. 2016): they have at least two small exons (< 50 bp) at the start of the gene that together code for a signal peptide, one large exon making up more than 70% of the coding sequence located near the end of the gene, and large numbers of serine-rich repeats (**Table 1**, **Figure 1**). Silk gland-specific expression was confirmed for five of the eight *A. luna* sericins (*ser1*, *serA-D*), based on a long-read transcriptome generated from silk glands extracted across different life stages (**Table 1**).

**Fig. 1:**
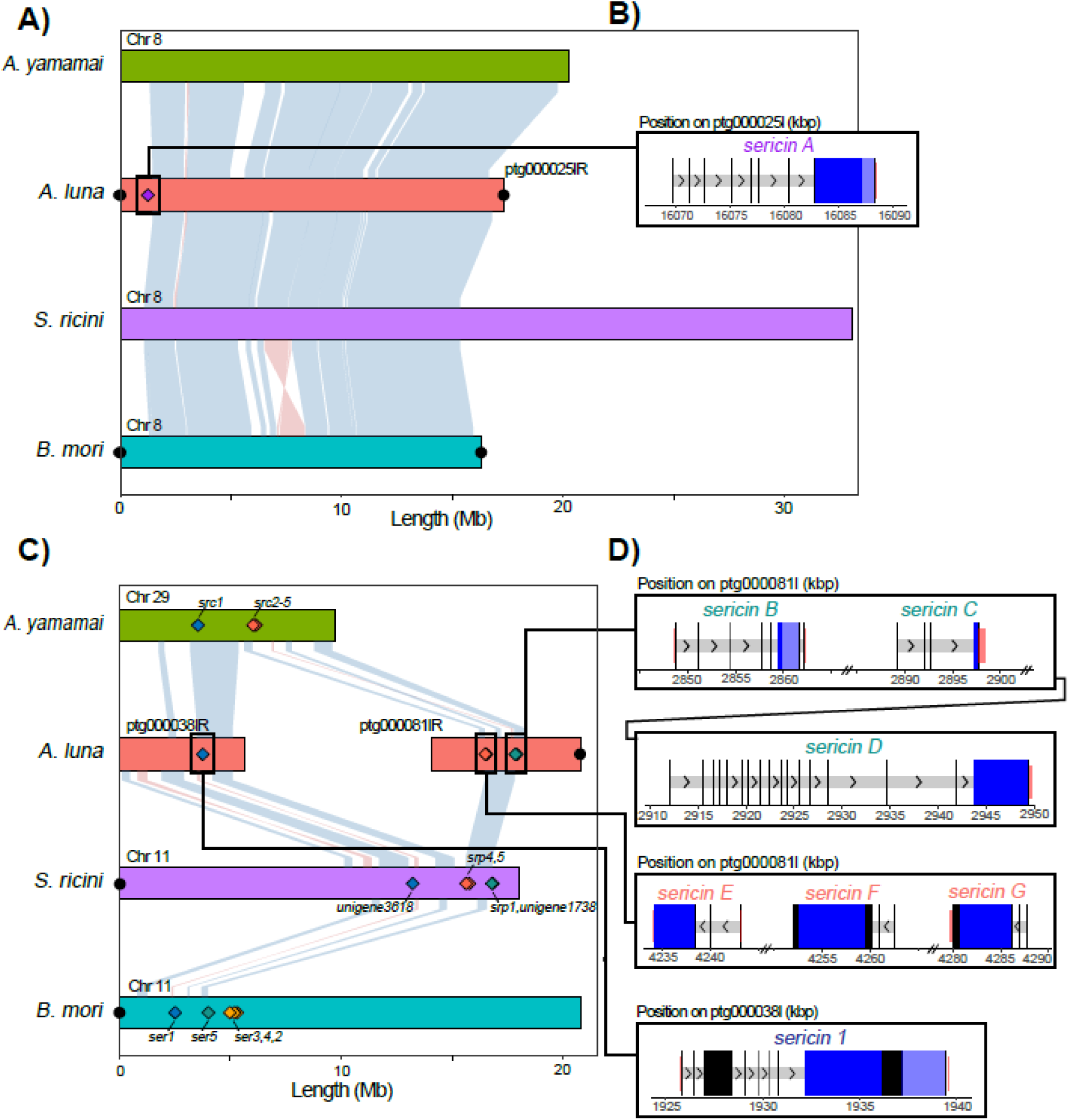
Comparison of sericin gene locations among four moth species. **(A,C)** BUSCO-derived ChromSyn synteny plots for *B. mori* chromosomes 8 **(A)** and 11 **(C)**. Light blue and light red lines connecting chromosomes represent synteny blocks of BUSCO genes, with blue indicating the same strand orientation and red indicating inversions. Contig names were retained from the respective NCBI assemblies; an “R” denotes contigs reversed in orientation. Filled black circles indicate predicted telomeres. Filled diamonds mark sericin gene locations, color-coded based on their genomic location. Names of *A. luna* sericins are listed in **(B)** and **(D)**, while sericin names for other species are listed on the plot. **(B,D)** Schematic diagram of *A. luna* sericin genes on ptg000025l **(B)** or ptg000038l and ptg000081l **(D)**, retaining the original chromosome coordinates. Wide black and blue boxes represent coding sequences (CDSs); introns are shown as narrow light gray boxes. Alternative exons (if present) are shown as wide dark gray boxes and untranslated terminal regions (UTRs) are marked in red. Primary and secondary repeat regions are shown in dark blue and light blue, respectively. Arrows in introns depict the direction of transcription. Chr = chromosome.

**Table 1.** Gene and protein characteristics for *A. luna* sericins. Silk gland expression was based on an IsoSeq silk gland transcriptome. Length, molecular weight, and Isoelectric point were extracted from encoded proteins after removal of the signal peptide in Geneious. R1 and R2 represent Repeat 1 and Repeat 2, as marked in Figure 1. GP = Genomic proximity; SS = Sequence similarity, SM = Sequence motifs, MW= Molecular weight, aa = amino acids, IP = isoelectric point

|  | SerA | Ser1 | SerB | SerC | SerD | SerE | SerF | SerG |
| --- | --- | --- | --- | --- | --- | --- | --- | --- |
| Identification method | SS, SM | SS, SM | GP | SS, SM | GP | SS, SM | SS, SM | SS, SM |
| Contig | ptg0000251 | ptg0000381 | ptg0000811 | ptg0000811 | ptg0000811 | ptg0000811 | ptg0000811 | ptg0000811 |
| Number of Exons | 8 | 8 | 7 | 4 | 19 | 3 | 3 | 3 |
| Protein length (aa) | 2012 | 3017 | 947 | 241 | 2439 | 2747 | 2082 | 1439 |
| Protein MW (kDa) | 190.67 | 297.64 | 93.31 | 23.82 | 248.03 | 264.77 | 201.81 | 140.64 |
| Protein IP | 2.99 | 6.07 | 3.87 | 2.96 | 3.26 | 7.12 | 5.20 | 5.24 |
| Repeat number | R1: 19<br>R2: 9 | R1: 13<br>R2: 14.5 | R1: 9<br>R2: 41 | 89 | 21 | 37 | 65 | 47 |
| Repeat length (aa) | R1: 64-76<br>R2: 41-45 | R1: 105-114<br>R2: 51 | R1: 13-15<br>R2: 15 | 21 | 8 | 38 | 38 | 38 |
| Silk gland expression | Yes | Yes | Yes | Yes | Yes | Unknown | Unknown | Unknown |

### Comparison of genomic locations of sericin genes in Saturniidae and *Bombyx mori*

To assess how sericins are related to each other, both within the same species and across species in the superfamily Bombycoidea, we evaluated whether sericin gene locations are conserved across species. A genome-wide synteny map of four different genomes (**Supplementary Figure S1**), including *A. luna*, two saturniid model species (*A. yamamai* and *S. ricini*), and the domesticated silkworm (*B. mori*), was generated using a set of highly conserved, single copy genes (Edwards et al. 2022). *Actias luna* sericin genes were located on three different contigs (**Table 1**, **Figure 1A,C**), but two of these (ptg000038l and ptg000081l) share synteny blocks of conserved genes with *A. yamamai* chromosome (Chr) 29, *S. ricini* Chr 11, and *B. mori* Chr 11 (**Figure 1C**, **Supplementary Figure S1**). Although chromosome rearrangements are frequent among the four species included in our synteny plot (*A. luna*, *A. yamamai*, *B. mori,* and *S. ricini*; **Supplementary Figure S1**), chromosome fission is rare in Lepidoptera (Wright et al. 2024). We thus hypothesize that *A. luna* contigs ptg00081l and ptg000038l represent one poorly assembled chromosome that contains seven out of the eight *A. luna* sericins.

Ptg000025l contains a single sericin gene, which we named *sericin A* (*serA*). *SerA* consists of seven short exons, followed by one large exon that contains two different repeat sequences (**Figure 1D**, **Table 1**). No previously characterized sericins were present in the orthologous chromosome 8 in *A. yamamai*, *S. ricini*, or *B. mori* (**Figure 1C**). *Sericin 1* (*ser1*), located on ptg000038l, is the largest *A. luna* sericin and contains eight different exons, two of which are alternative exons that are not always present (**Table 1**, **Figure 1D**). Its longest exon contains two different sets of repeats (**Table 1**, **Figure 1D**). Its genomic location is highly conserved in all four species included in the synteny plot (**Figure 1C**), and possibly within the superfamily Bombycoidea.

The other six sericins (*serB-G*) are found in a 1.5 Mb-long region on ptg000081, evenly spread in two distinct gene clusters (**Figure 1**). The first of these gene clusters, spread across almost 105 kb, contains *serB-D* (**Figure 1**). These three sericins vary strongly in the number of exons (4-19), repeat number (21-89), and total length (241-1439 amino acids) (**Table 1**, **Figure 1D**). The same genomic region also contained sericins in *S. ricini* (*Srp1*, *Unigene1738*) and *B. mori* (*ser5*), but lacked known sericins in *A. yamamai* (**Figure 1C**). The second gene cluster, consisting of *serE-G*, covers an even smaller genomic region of under 55 kbp. *SerE-G* all consist of a three exon-structure and a 38 amino acid repeat in their longest exon (**Table 1**, **Figure 1D**). We identified sericins in the same genomic region for the two other saturniid species, *A. yamamai* (*Src2-5*) and *S. ricini* (*srp4,5*), but not for *B. mori* (**Figure 1A**). The remaining *B. mori* sericins, *ser2-4*, were in a more distant region on the same chromosome, which lacked sericins in any of the three included saturniid species (**Figure 1C**).

### Grouping of sericins based on protein similarities

To further characterize relationships between saturniid sericins, a phylogenetic network was generated based on pairwise protein identities, including all translated *A. luna* sericins and the translated sericin sequences from the sericin dataset described under “Identification of sericins in the *Actias luna* genome” (**Supplementary Table S1**). We recognize four different groups of sericins in the family Saturniidae, based on sequence similarity at the protein level among eight saturniid moths (*A. assamensis*, *A. luna*, *A. pernyi*, *A. selene*, *A. yamamai*, *H. cecropia*, *R. newara*, and *S. ricini*) and *B. mori* (**Figure 2**).

**Fig. 2:**
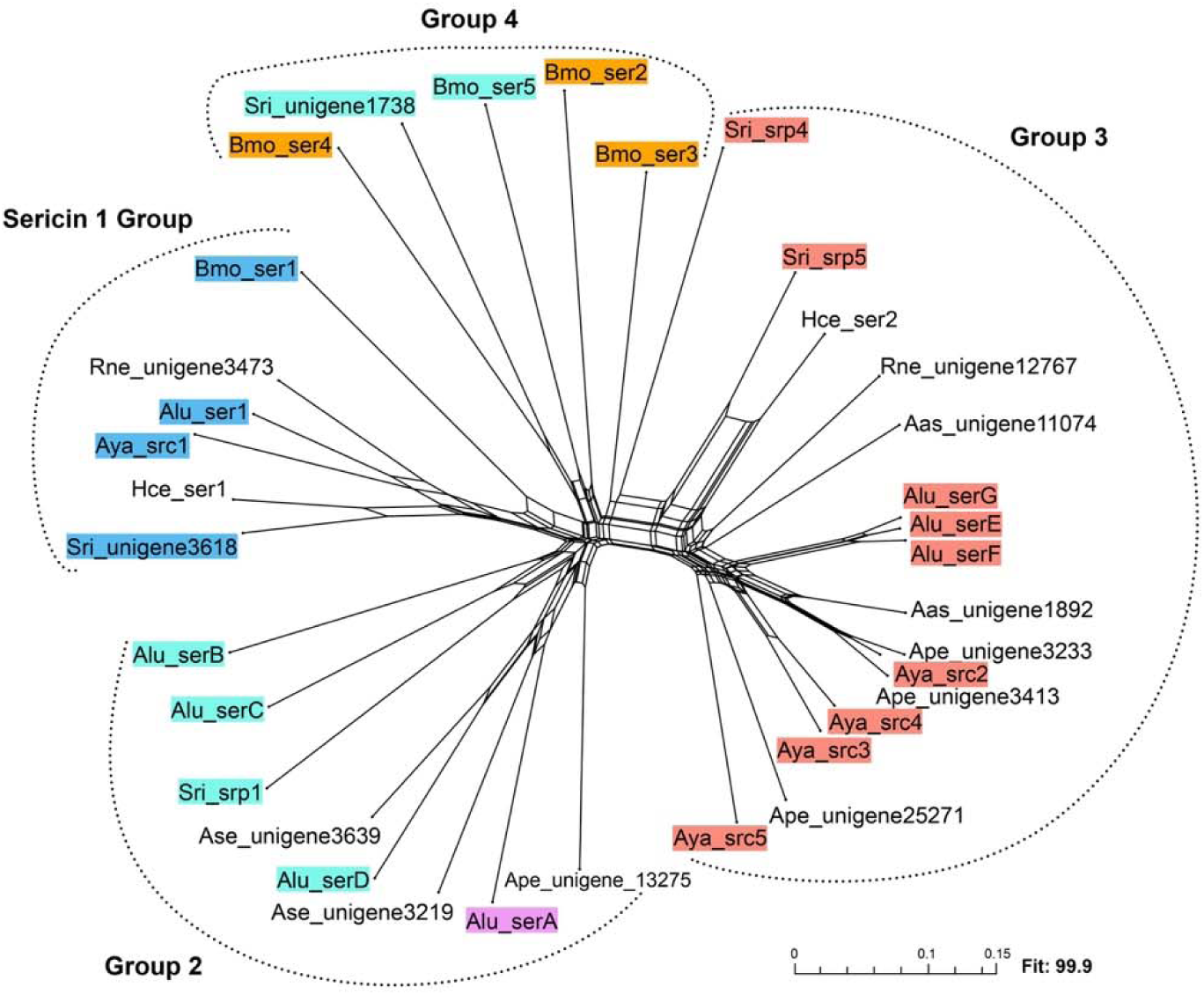
Sericin phylogenetic network. A phylogenetic network based on pairwise distances between sericin protein sequences was generated with SplitsTree. Sericins were color-matched with colors used in Figure 1 where applicable. Sericin groups are represented by dotted lines. Abbreviations: Aas: *Antheraea assamensis*, Alu: *Actias luna*, Ase: *Actias selene*. Ape: *Antheraea pernyi*, Aya: *Antheraea yamamai*, Hce: *Hyalophora cecropia*, Bmo: *Bombyx mori*, Rne: *Rhodinia newara*, Sri: *Samia ricini*, srp: serine-rich protein.

The first group contains six sericins, each from a different species (**Figure 2**). These include *B. mori* ser1 and three sericin proteins encoded by genes located in the same genomic location as *B. mori ser1* in other species (*Alu_ser1*, *Aya_src1* and *Sri_unigene3618*; **Figure 1**). Each sericin protein within this cluster contains a conserved CXCX motif at the N terminus (**Supplementary Figure S2**), representative of sericin 1 sequences across Lepidoptera (Wu et al. 2022). We thus named this group the Sericin 1 Group (Figure 2). The second group comprises eight different sericin sequences, belonging to only four of the nine analyzed species with a notable absence of *B. mori*. Due to the absence of *B. mori* sequences, we have separately grouped these proteins and named it Group 2 (Figure 2). The four *A. luna* sericin genes (*serA-D*) encoding Group 2 proteins are found in two separate genomic regions: *serB–D* are closely clustered, while *serA* resides on a different chromosome (**Figure 1A,C**). Interestingly, despite their genes being located on different contigs, serD is more closely related to serA than to serB or serC, which are part of the same gene cluster (**Figure 1**, **Figure 2**). Homologs of serA and serD are also found in *A. selene*, but no closely related sequences were identified outside the genus *Actias*.

The third group is well-represented in almost every saturniid taxon included in our study but does not contain *B. mori* sericin sequences (**Figure 2**). As such, we refer to this group as Group 3. The *A. luna* sericins in this group (SerE-G) are highly similar at the protein level (**Figure 2**), despite considerable variation in the number of repeats (**Table 1**). The *A. yamamai* genome also shows an expansion of Group 3, with at least four distinct members (*Src2-5*) (Žurovec et al. 2016) that are less conserved (**Figure 2**). All sericins in Group 3 share a conserved 38-amino acid repeat motif (**Table 1**) (Žurovec et al. 2016). Although two of the *S. ricini* sericins, serine-rich protein 4 and 5 (srp4, srp5), appear to be only distantly related to other Group 3 sequences (**Figure 2**), they are found in the same genomic region (**Figure 1**) and contain a 38-amino acid repeat motif, found in all Group 3 sericins. As such, we included these sericins in Group 3. Interestingly, *Samia ricini* srp4 and srp5 share some similarity with *B. mori* ser3 (**Figure 2**). However, they exhibit different repeat lengths and the gene coding for *B. mori* ser3 is in a different genomic region than *S. ricini srp4* and *srp5* (**Figure 2**). Thus, the relationship between *B. mori* sericin 3 and Group 3 sericins remains unresolved. The remaining *B. mori* sericins (ser2-5) form a separate fourth group, that includes one *S. ricini* sericin (**Figure 2**). Notably, the genes coding for *B. mori* ser5 and *S. ricini* unigene1738 share a genomic location with saturniid genes encoding Group 2 sericins (**Figure 1**), but the proteins share greater sequence similarity with *B. mori* ser2 and ser4. This final group, which lacks any *A. luna* sericins, is referred to as Group 4.

### Differential expression of *Actias luna* sericins across life stages

Previously generated RNA-sequencing data (Markee et al. 2024) were used to examine whether sericin gene expression varies between life stages. We compared first, fourth, and last (fifth) instar caterpillars. Although the data are not silk gland-specific, sericin gene expression outside of the silk glands is very low (Dong et al. 2019; Wu et al. 2024) or undetectable (Guo et al. 2022; Wu et al. 2022; Žurovec et al. 2016). We observed two distinct expression patterns among *A. luna* sericin genes. Group 2 sericins (**Figure 2**; *serA-D*) exhibited high expression levels in at least one of the two earlier life stages and their expression levels in the last instar (L5) dropped to close to zero. In contrast, *ser1* and the Group 3 sericins (**Figure 2**; *serE-G*) had low expression levels across life stages and showed a trend towards higher expression levels in the last instar (L5). Specifically, each Group 2 sericin (**Figure 2**; *serA-D*) exhibited significantly higher expression levels in the first instar (L1) compared to the last instar (L5) (**Figure 3**). For *serB* and *serD*, expression levels declined markedly from the first to fourth instar. In contrast, *serA* showed peak expression in the fourth instar and *serC* exhibited a similar upward trend (**Figure 3**). The expression levels of *serG* were negligible in each sample, but *ser1*, *serE*, and *serG* reached their highest expression levels in a L5 caterpillar that also exhibited high expression levels for *FibH* (**Figure 3**). Of these, *serE* was the only one for which the gene expression level in L5 was significantly elevated compared with L1.

**Fig. 3.**
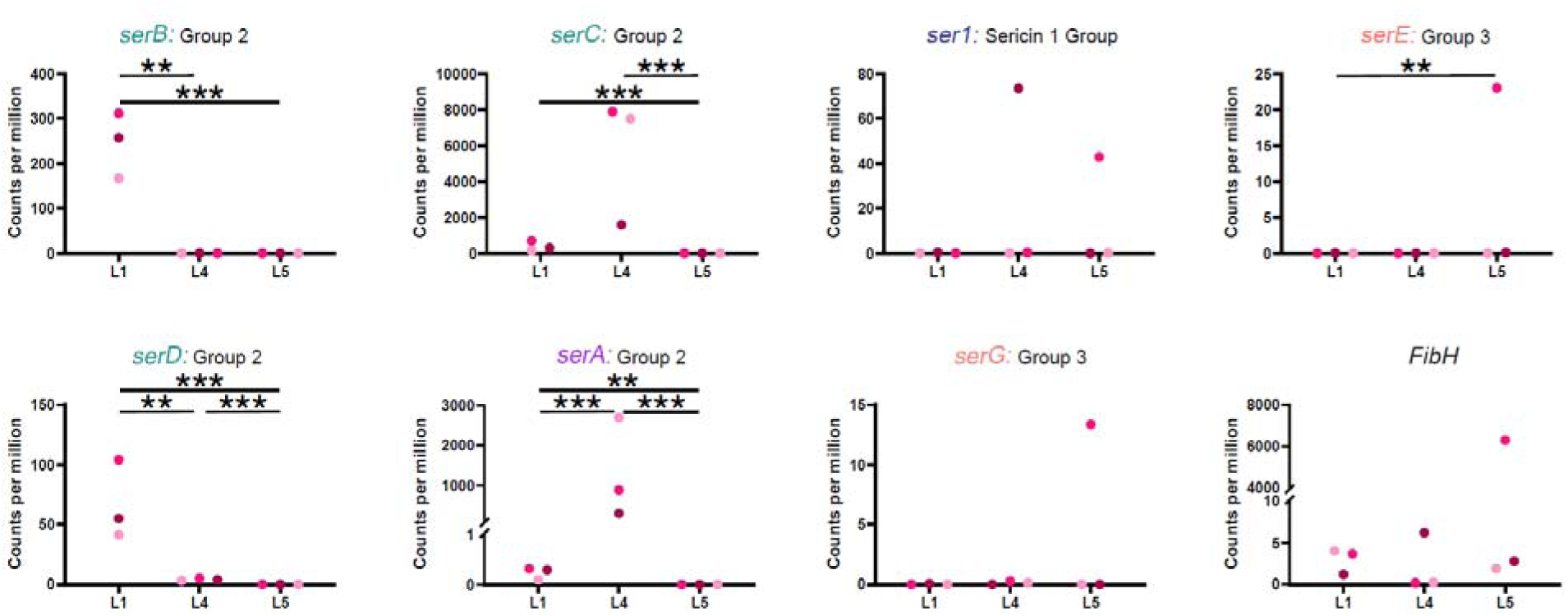
Sericin and *FibH* gene expression during Luna moth development. Read counts were obtained by RNA sequencing of the whole body (first instar; L1) or the abdomen (fourth instar; L4 and fifth instar; L5) and were normalized in edgeR (Robinson et al. 2010). Expression levels of *serF* were negligible and are not shown here. Sericin names were color-matched with **Figure** and group names were adapted from Figure 2. The *FibH*-gene was previously identified by Markee et al. (2024). Significance levels are denoted by *, ** and *** (respectively *p* < 0.05, *p* < 0.01 and *p* < 0.001).

### Protein composition and repeat motifs across *Actias luna* sericins

To explore potential functional differences among sericins, we compared the amino acid composition and the repeat sequences for the eight predicted *A. luna* sericin proteins. Most repeats were rich in serine and threonine (**Figure 4**), which respectively comprised 22–31.2% and 14.9–34.6% of total residues. In contrast, in serA, serine accounted for 59% due to extended serine stretches, but a corresponding decrease in threonine balanced the overall serine/threonine ratio and kept it comparable to the other *A. luna* sericins. Based on a combination of their amino acid proportions and their expression levels across life stages, *A. luna* sericins can be functionally separated. Group 2 sericins (serA-D; **Figure 2**), that exhibit high expression levels at earlier instars (**Figure 3**), contain high levels of acidic residues (aspartic acid: 5.8-22%, glutamic acid: 2.1-5.6%) but low levels of basic amino acids (combined: 1.6-3.8%). Their repeats contain long stretches of serine and threonine residues, interspersed with aspartic acid, glutamic acid, alanine, and, to a lesser extent, valine residues (**Figure 4A**). SerB additionally exhibited elevated levels of proline. In contrast, the remaining sericins – ser1 and the three Group 3 sericins (serE-G) – reach their highest expression levels in the final instars and contain high levels of basic residues (> 6.5%), glycine (13.2-20.5%), and uncharged polar residues (9.0-17.3%), but low levels of acidic residues (5.9-7.4%). *Actias luna* ser1 is defined by two sets of repeats, both of which are proline-rich and contain a recurring motif consisting of a glycine followed by five serine or threonine residues. The first repeat contains two stretches that are high in histidine, glycine, and proline residues, while the second repeat is high in tyrosine, aspartic acid, and arginine (**Figure 4A**). Finally, the three Group 3 sericins (serE-G) contain high levels of tyrosine, particularly within a recurring S/TYTS motif (**Figure 4A**). The differences in basic and acidic residues among *A. luna* sericins are also reflected in their pI, which is the lowest for Group 2 sericins (**Table 1**).

**Fig. 4.**
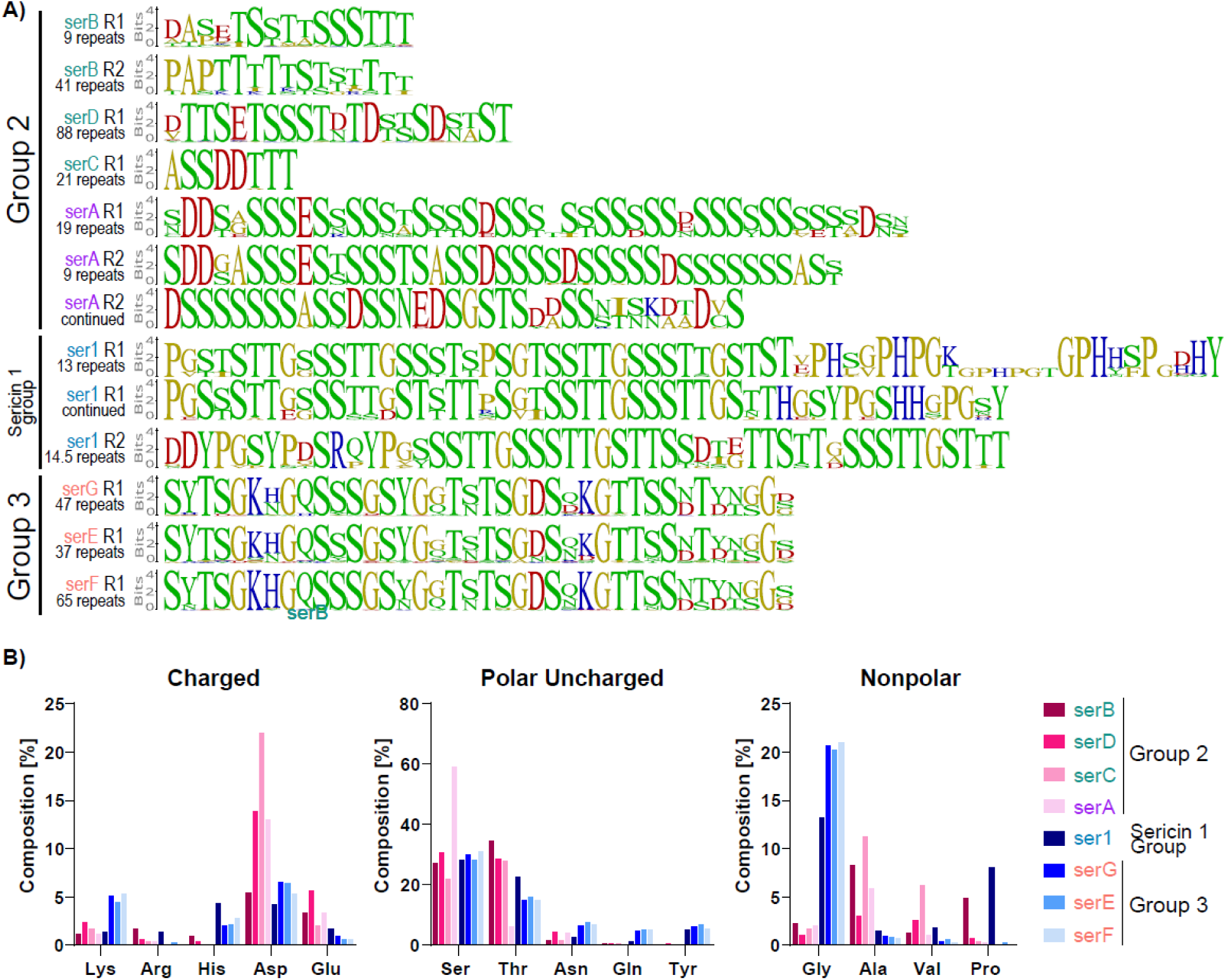
Protein composition of *A. luna* sericins. **(A)** Repeat sequence motifs for each *A. luna* sericin, separated per repeat where applicable. Amino acids are colored based on their polarity (blue: basic, red: acidic, green: polar, yellow: non-polar). Sericins are ordered based on when they reached the highest expression level (Figure 3), their names were color-matched with Figure 1, an groupings based on Figure 2 were marked in the legend. The number of repeats is listed below the name and repeat number. **(B)** Amino acid composition of *A. luna* sericins. Only abundant amino acids or amino acids that strongly varied between sericins are shown. Sericins were ordered and colored as in (A).

## Discussion

### The *Actias luna* genome contains at least eight putative sericins

Our study identifies *A. luna* as one of the species with the highest number of characterized sericins to date, surpassed only by *Galleria mellonella* (Pyralidae) for which at least 12 sericin-like proteins were identified (Wu et al. 2022). Each of the eight sericins identified here matches structural characteristics that appear to be a defining feature of this gene family (Dong et al. 2019; Guo et al. 2022; Kludkiewicz et al. 2009; Takasu et al. 2007; Wu et al. 2024; Žurovec et al. 2016): two short initial exons that together encode the signal peptide and one long exon that contains a large number of serine-rich repeats (**Figure 1**). Additionally, each sericin is either expressed in our long read, silk gland-specific transcriptome (*ser1*, *serA-D*, **Table 1**), or orthologous to genes found in other species to be major components of cocoon silk – *serE-G* are orthologues of *A. yamamai Srn2-5* and *H. cecropia src2* (**Figure 1,2**)(Rouhová et al. 2024; Žurovec et al. 2016). The absence of the *serE-G* in our silk gland transcriptome is likely due to limited sampling – only one individual per life stage was included, without specifically targeting a prepupal caterpillar – and the low quality of our sequencing data, resulting in a substantial loss of raw reads during quality filtering steps (**Supplementary Table S2**). The lower number of reported sericins in the seven other saturniid moth species is likely due to incomplete gene characterization rather than true absence of these genes. For instance, most studies identifying sericins focus primarily on cocoon silk and/or late larval instars, potentially missing sericins expressed during earlier larval instars (Dong et al. 2015; Rouhová et al. 2024; Tsubota et al. 2016; Žurovec et al. 2016).

### Gene duplications led to the origin of four different sericin groups

Due to their high repeat content (**Figure 1**), the resulting fast evolutionary rates (Babcock et al. 2003; Cheng & Chen 1999; Delihas 2011; King 2024), and the highly biased amino acid content (**Figure 4**), assessing evolutionary relationships among sericins remains a challenge (Kmet et al. 2023; Tsubota et al. 2021; Wu et al. 2022). Based on genomic locations and pairwise sequence similarities at the protein level, we were able to separate the saturniid sericins into four groups (**Figures 1, 2**). Although genomic locations (**Figure 1**) and protein similarities (**Figure 2**) are generally congruent among the sericins included in our study, there are a few discrepancies. For instance, *B. mori ser5* and *S. ricini unigene1738*, both placed in Group 4 based on protein level similarities (**Figure 2**), share their genomic locations with *A. luna serB-D* of Group 2 (**Figure 1C**). Similarly, *B. mori* ser3 and Group3 sericins are not well separated based on protein sequences (**Figure 2**) but differ in genomic location (**Figure 1C**) and repeat lengths. Increased sampling of sericins is required to better understand the relationships between the different sericin groups described in the current study. Of the four sericin groups identified in this study, the Sericin1 Group is the only one that is shared between *A. luna* and *B. mori*, while the others appear to exhibit no or minimal overlap between Saturniidae and Bombycidae (**Figures 1,2**). As silk production in *B. mori* has been the subject of an incredibly large number of studies on gene expression, proteomics, and transcriptomics (e.g. (Dong et al. 2013; Guo et al. 2023; Masuoka et al. 2024; Peng et al. 2019; Zhang et al. 2015), we consider it unlikely that the *B. mori* genome contains additional unidentified, functional sericins (but see Wu et al. (2024) for a recently identified sericin-like gene). Similarly, we were unable to identify any *A. luna* sericins closely related to *B. mori ser2-5* or *S. ricini unigene1738*. Our data thus suggests that the sericins in *B. mori* and *A. luna* exhibited distinct evolutionary trajectories, with an expansion of Group 4 in the lineage of *B. mori* and expansions of Group 2 and Group 3 in Saturniidae.

Sericin Group 2 and Group 3 are represented in *A. luna* by four and three sericins, respectively. While *serA* of Group 2 is isolated on ptg000025l – the *A. luna* homologue to *B. mori* Chr8 -, *serB-D* of Group 2 and *serE-G* of Group 3 are in two gene clusters within relative genomic proximity of each other on ptg000081l – homologous to *B. mori* chr11 (**Figure 1**). The close genomic proximity and high sequence similarity within each of these clusters suggests that they both reflect tandem duplications (**Figure 1, 2**). In particular, *serE-G* are nearly identical based on overall gene structure (**Figure 1B**), sequence similarity (**Figure 2**), gene expression or rather the absence thereof (**Figure 3**), repeat motifs (**Figure 4A**) and amino acid content (**Figure 4B**), and only clearly differ in their repeat number (**Table 1**, **Figure 1B**) and the sequence of their introns (**Supplementary Figure S3**). Near-identical gene copies are often thought to represent very recent duplications, and are expected to either diverge or disappear (Lynch & Conery 2000). Alternatively, gene conversion, a process where homologous recombination results in the unidirectional transfer of genetic material from a donor gene to an acceptor gene, can also result in near-identical gene copies (Chen et al. 2007). The high level of divergence among the introns, and to a lesser extent the non-repetitive C-terminus, of *serE* compared to *serF* and *serG* (**Supplementary Figure S3**) suggests that gene conversion could explain the similarity of their repeat sequences (**Figure 4A**). Recently duplicated genes often experience high levels of positive selection and undergo rapid functional divergence, either at the sequence level (Lynch & Conery 2000; Qiao et al. 2019) or at the expression level (Brasó-Vives et al. 2022; Cai & Des Marais 2024; Huerta-Cepas et al. 2011). In rare instances, sequence similarities are retained when increased gene dosage is beneficial to the organism (Hahn 2009; Perry et al. 2007; Sackton et al. 2007) or through hypofunctionalization (Birchler & Yang 2022; Brasó-Vives et al. 2022), a process where the expression of each copy decreases in order to retain final expression levels at the pre-duplication level. The Group 3 sericins in other Saturniidae are also organized in a single gene cluster (e.g. *A. yamamai Src2-5*, *S. ricini srp4,5*; **Figure 1C**), but they diverged more than in *A. luna* (**Figure 2**). Although the lineage-specific expansions of Group 3 sericins could contribute to increased dosage, we hypothesize that the high similarities among *A. luna serE-G* reflects their recent origin or gene conversion and that they will eventually functionally diverge.

The other *A. luna* sericin gene cluster (Group 2 sericins *serB-D*) is more divergent at the protein level, suggesting it is the result of an older gene duplication. Surprisingly, *A. luna* s*erC* seems to be more closely related to *A. luna serA*, located on a different chromosome, than to the two sericins in its own gene cluster (*serB, D*) (**Figure 1, 2**). *SerA* might represent a relatively recent across-chromosome duplication of *serC*. Orthologues of both *serA* and *serC* are also present in *A. selene*, suggesting such duplication would have happened in a common ancestor of *A. luna* and *A. selene* (**Figure 2**). *A. luna* s*erA* is only the second confirmed case of a sericin found by itself on a chromosome, while most sericins are found within the same chromosome, homologous to *B. mori* Chr11 (Guo et al. 2022; Wu et al. 2022; Žurovec et al. 2016). The only other sericin located outside this chromosome is *sericin P150* (Kludkiewicz et al. 2019; Wu et al. 2024), which is unrelated to serA because it differs in its chromosomal location (**Supplementary Figure S1**) and in the presence of a conserved CXCXCX-motif in its N-terminus that is absent in *A. luna* serA (Kludkiewicz et al. 2019; Wu et al. 2024).

The evolution of sericins might be driven by two main factors. First, the high repeat content of sericins and their close genomic proximity (Dong et al. 2019; Wu et al. 2022; Žurovec et al. 2016) allows for increased gene duplication rates, generating redundant gene copies on which evolution has free play. Notably, recent expansions of tandemly duplicated genes have repeatedly been shown to play an important role during species differentiation and adaptation (Brown et al. 1998; Clifton et al. 2017; Delihas 2020; Jugulam et al. 2014; Newcomb et al. 2005; Perry et al. 2007). Second, evolutionary rates can be further accelerated by the high intragenic repeat content of sericins. Intragenic, repeat-mediated duplications can alter gene structure and sequence (Babcock et al. 2003; Cheng & Chen 1999; Delihas 2011; King 2024). Additionally, intragenic nonreciprocal recombination (gene conversion) or unequal crossing over among intragenic repeats can lead to fast homogenization of the repeat region and can quickly spread new mutations across a whole gene (Garb et al. 2007; Hayashi & Lewis 2000). The formation of gene clusters and the resulting lineage-specific expansions and deletions of sericins appears to be a hallmark of sericin evolution, as this phenomenon has also been reported in the lepidopteran family Pyralidae (Žurovec et al. 2016).

### *A. luna* sericins exhibit functional specialization

Gene duplications can facilitate neofunctionalization or subfunctionalization of the redundant gene copies, as duplicated genes are free to accumulate mutations without disrupting the ancestral gene function (Birchler & Yang 2022; Kuzmin et al. 2022). In this study, we hypothesize that *A. luna* sericins have undergone subfunctionalization, with different paralogs exhibiting similar functions at different life stages.

The four *A. luna* Group 2 sericins (*serA-D*, **Figure 2**) are expressed in the first and/or fourth instar but exhibit low to undetectable levels in the final instar (**Figure 3**), and we thus refer to them as larval sericins (**Figure 5**). The remaining *A. luna* sericins (*ser1*, and Group 3 sericins *serE, serF* and *serG*, **Figure 2**) exhibit overall low expression levels in our study, but reach their highest expression levels in a last instar individual that also exhibits increased levels of *FibH* expression (**Figure 3**). Although behavioral data were not recorded for the dissected caterpillars, this suggests that this caterpillar was actively spinning silk for cocoon creation at the time of dissection. If so, *A. luna ser1* and the *A. luna* Group 3 sericins, *serE*-*G,* would represent cocoon sericins (**Figure 5**). In support of this idea, the proteins coded by their orthologues in *A. yamamai* (*Srn1-5*), *H. cecropia* (*src1, src2*), and *B. mori* (*ser1*) (**Figure 1, 2**) are all major components of cocoon silk (Peng et al. 2019; Rouhová et al. 2024; Takasu et al. 2002; Wu et al. 2024; Žurovec et al. 2016). Intriguingly, we observed striking differences in amino acid composition between *A. luna* cocoon sericins (ser1 and Group 2 sericins serE-G) and larval sericins (Group 3; serA-D), with the latter having higher levels of charged residues and lower tyrosine levels in *A. luna* cocoon sericins. In further support of sericin differentiation across life stages (**Figure 5**), *A. luna* silk fibers from different life stages exhibit compositional differences in the sericin-containing outer coating, but not in the inner fibroin core (Eccles et al. *Under Review*).

**Fig. 5.**
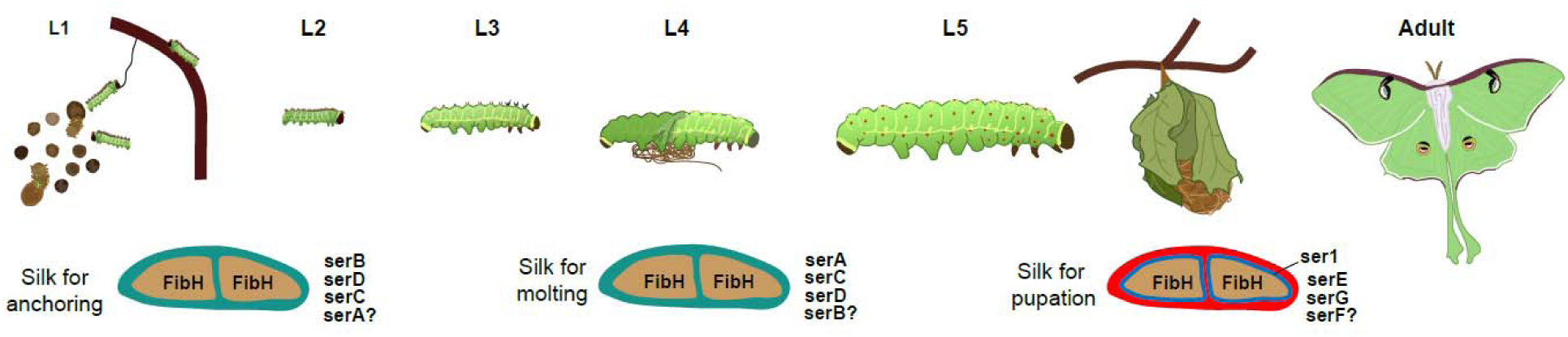
Proposed hypothesis on *Actias luna* silk sericin composition across different life stages. This scenario is based on our current findings and previously published work (see “*A. luna* sericins exhibit functional specialization”)

These observations are reminiscent of the well-studied *B. mori*, where differential expression of sericins was shown to drive functional divergence between larval and cocoon silks, with each being adapted to the caterpillar’s ecological needs (Guo et al. 2022, 2025; Kludkiewicz et al. 2009; Takasu et al. 2010). In *B. mori*, larval sericins (*B. mori* ser2, ser4, ser5 and serP150) also exhibit higher levels of charged amino acids (respectively, > 30% and < 20% of total amino acids) than cocoon sericins (*B. mori* ser1 and ser3) (Dong et al. 2013, 2019; Guo et al. 2023, 2022; Kludkiewicz et al. 2009; Peng et al. 2019; Takasu et al. 2002, 2007, 2010; Wu et al. 2024; Zhang et al. 2015) and the cocoon sericin *B. mori* ser1 has similarly elevated tyrosine levels as *A. luna* cocoon sericins. Additionally, *B. mori* larval sericins have lower serine levels than cocoon sericins (respectively, < 16% and >34%) (Dong et al. 2019; Guo et al. 2022; Takasu et al. 2010). The highly charged, low-serine larval sericins give *B. mori* silk a high adhesion to their host plant and substrate (Dong et al. 2019; Guo et al. 2022; Kludkiewicz et al. 2009; Takasu et al. 2010) and high rigidity and strength due to increased β-sheet proportion in early life stages (Guo et al. 2022; Peng et al. 2019) when compared to cocoon silk. The function of tyrosine residues in cocoon sericins is currently unknown, but the strong hydrogen bonds formed by tyrosine residues in *B. mori* silk fibroin aid in self-assembly (Partlow et al. 2016) and might similarly assist sericin assembly or interactions with other proteins.

This suggests that *A. luna* sericins could similarly drive the functions of the silk fiber outer coating. Serine levels in *A. luna* sericins are consistently high (>20%) without differences between larval and cocoon sericins as in *B. mori*, and the differences in charged residues are not as extreme. We hypothesize that the compositional differences between both species represent differences in their respective ecology, for instance in cocoon structure (Chen et al. 2012b). This high serine-content in *A. luna* sericins, observed across Saturniidae (Rouhová et al. 2024; Žurovec et al. 2016), is at odds with other investigated lepidopteran species, where serine content can drop as low as 12% in (Dong et al. 2019; Guo et al. 2022; Takasu et al. 2010; Wu et al. 2022). Additionally, *A. luna* ser1, and to a lesser extent serB, are unique among *A. luna* sericins because they exhibit significant proline levels. In spidroins, the major silk proteins of spiders, high proline levels are linked to increased silk elasticity (Savage & Gosline 2008), suggesting the high proline levels in *A. luna* sericin 1 might similarly increase silk elasticity. Life-stage specific sericin expression has been observed in a variety of lepidopteran taxa (Kludkiewicz et al. 2009; Masuoka et al. 2024; Peng et al. 2019; Takasu et al. 2002; Wu et al. 2024; Žurovec et al. 2013), and similar compositional trends might thus be found across Lepidoptera.

In addition to the life stage-specific regulation of sericin genes, they are also spatially regulated. Of the two main cocoon sericins in *B. mori*, *ser1* is expressed exclusively in the posterior end of the middle silk gland and the encoded protein is deposited as an inner sericin layer of *B. mori* cocoon silk, while *ser3* is expressed throughout the middle silk gland and its encoded protein forms an outer sericin layer (Dong et al. 2013; Guo et al. 2023; Takasu et al. 2010). Ser1 appears to be essential for successful silk assembly (Takasu et al. 2017) and its sequence and expression pattern are conserved across Lepidoptera (Guo et al. 2025; Kludkiewicz et al. 2019; Rouhová et al. 2021; Takasu et al. 2007) including Saturniidae: *ser1*-orthologues in two saturniid taxa – *A. yamamai Srn1* and *H. cecropia Src1* (**Figure 1,2, Supplementary Figure S2**) – are only expressed in the posterior part of the middle silk gland (Rouhová et al. 2024; Žurovec et al. 2016). *Antheraea yamamai* and *H. cecropia* do not have a direct orthologue of *B. mori ser3* (**Figure 1,2**) and instead express Group 3 sericins (respectively *Srn2-5* and *Src2;* **Figure 1,2**) throughout the middle silk gland of prepupal caterpillars (Rouhová et al. 2024; Žurovec et al. 2016). Thus, the two-layered structure of the sericin outer coating might be retained in Saturniidae, with the inner ser1 layer being highly conserved and saturniid Group 3 sericins (including *A. luna serE-G*, **Figure 1,2**) replacing *B. mori* ser3 in the outer layer (**Figure 5**). This hypothesis is further supported by the elevated glutamine levels found in both *A. luna* Group 3 sericins and *B. mori* ser3, which could play a role in increasing sericin adhesion (Dong et al. 2019). While *B. mori* larval silks exhibit a similar multi-layered sericin organization (Dong et al. 2019), silk gland region-specific data for larval sericins are currently lacking for Saturniidae and it is thus unclear whether larval silks in Saturniidae exhibit a similar multi-layered sericin composition.

The similar patterns of sericin temporal expression, spatial expression, and composition between Saturniidae and *B. mori* is particularly surprising considering that their respective sericins represent mostly separate gene expansions (**Figure 1,2**). In *B. mori*, Group 4 sericins diverged into larval and cocoon sericins with distinct expression patterns and amino acid compositions. In *A. luna* and other Saturniidae, Group 2 and Group 3 sericins have convergently specialized into larval and cocoon sericins, respectively. Sericin 1, the only sericin for which we found no duplications, seems to be conserved across the included lineages, and likely preserved its function. Further investigations outside Bombycidae and Saturniidae are needed to establish which represents the ancestral situation, and whether other lineages of Lepidoptera convergently evolved multiple sericins specialized for particular life stages.

### Concluding remarks

Most genetic studies and genotype-phenotype analyses in lepidopteran silk research have focused on *FibH* and its encoded protein - the primary component of the silk fiber core in Lepidoptera and Trichoptera and the largest silk protein in both insect orders (Heckenhauer et al. 2023; Yonemura et al. 2009; Zhang et al. 2024). However, sericins form the bulk of the outer silk fiber coating (Dong et al. 2013; Guo et al. 2022; Rouhová et al. 2024) and can influence silk viscosity (Peng et al. 2019), adhesion (Guo et al. 2022; Lee et al. 2018), and self-assembly (Takasu et al. 2017). In this study, we demonstrate that duplications of sericins, a family of highly repetitive silk proteins specific to lepidopterans, enable dynamic shifts in silk composition, potentially reflecting adaptation to changing ecological and functional demands. Sericins are characterized by a high repeat content (**Table 1**, **Figure 1B**, **Figure 4A**), frequent organization into clusters of closely related paralogs (**Figure 1**) (Dong et al. 2019; Wu et al. 2022; Žurovec et al. 2016), and substantial variation in repeat number and sequence composition (**Table 1**, **Figure 4A**). As such, they represent a compelling system for studying the dynamics and evolutionary consequences of gene duplication. Our findings suggest that sericins have undergone extensive gene expansions and deletions in Saturniidae and Bombycidae, facilitating both subfunctionalization and convergent evolution across taxa. While fibroin proteins are also known for their evolutionary variability - even displaying divergent alleles within a single caterpillar (Frandsen et al. 2023; Markee et al. 2024) - the frequent duplication of sericin genes, followed by changes in expression patterns and repeat content, may allow the silk coating to evolve even more rapidly.

## Methods

### Caterpillar Rearing

All *A. luna* caterpillars descended from wild-caught females collected in the southeastern United States in 2021. Eggs were collected in a paper bag and transferred to a US Department of Agriculture (USDA) containment lab at the University of Florida’s McGuire Center for Lepidoptera and Biodiversity. After hatching, caterpillars were kept together in clear plastic cups with lids (16 oz.) and were fed with fresh American sweetgum (*Liquidambar styraciflua*).

### Short-read RNA Sequencing and Read Alignment

Samples for Illumina short-read RNA sequencing were part of a larger study described previously (**Supplementary Table S3**) (Markee et al. 2024). Reads from 28 samples, originating in a variety of life stages and body parts, were trimmed based on their quality with trimmomatic v0.39 (removing bases with a score under 25 from either end, as well as using a sliding window of 3 bases with a minimum average quality of 20) (Bolger et al. 2014). Additionally, trimmomatic was used to remove adapter sequences, using the “Truseq3-PE-2.fa” file provided by trimmomatic v0.39. Pairs for which both reads had a final length of over 50 were retained for further analysis. Sequencing errors were corrected using Rcorrector v 1.0.4 (Song & Florea 2015) and unfixable reads were removed using a publicly available python script (rm_rcorrector_unfixable.sh, https://github.com/harvardinformatics/TranscriptomeAssemblyTools). Filtered reads were aligned to the genome of *A. luna* (GenBank accession number GCA_039707435.1) (Markee et al. 2024), using Subread v2.0.6 (Liao et al. 2013), allowing multimapping and using a genome feature file (gff) generated by Markee et al. (2024) that included sericins and was adapted for use by subread. The number of reads mapping to each gene was counted with featureCounts (Liao et al. 2014), assigning fractional counts to multimapping reads. Raw read counts were subsequently imported into R v4.2.2 (R Core Team 2010) and normalized with the counts per million method using edgeR v3.40.2 (Robinson et al. 2010) for plotting purposes. For differential expression analysis in edgeR, raw reads were normalized using the TMM method (Robinson et al. 2010) and a quasi-likelihood F (QLF) test was used to assess differences between selected groups (whole body 1^st^ instar, abdomen 4^th^ instar, abdomen 5^th^ instar; n=3 for each group), using the Benjamini-Hochberg method to correct for multiple testing.

### Long-read Silk Gland Transcriptome

Silk glands were carefully extracted from a single individual at each representative life stage (instars 1-5 – L1-5) by decapitating caterpillars live over ice while submerged in Qiagen RNAprotect Tissue Reagent (Qiagen, Cat #76104). Glands were immediately flash frozen in liquid nitrogen in 1.8 mL microcentrifuge tubes and stored at -80°C. RNA extractions, library preparation, sequencing and initial subread processing were performed at the University of Florida’s Interdisciplinary Center for Biotechnology Research (RRID:SCR_019152). RNA was isolated using a Qiagen RNeasy micro extraction kit (Qiagen, Cat #74004) following manufacturer protocol and assessed for concentration and quality using a Qubit fluorometer and the Agilent 2100 Bioanalyzer (Agilent Technologies, Inc). SMRT Bell IsoSeq libraries were prepared for PacBio SEQUEL IIe according to manufacturer protocol (Pacific Biosciences, Cat #PN 101-763-800) with few modifications. RNA preps were cleaned and concentrated using the ZYMO Research RNA Clean and Concentrator kit (ZYMO Research, Cat #R1015). Total RNA from each larval instar silk gland was diluted to approximately 100 ng, after which full-length cDNA was synthesized with amplification as described in the protocol for Low Input RNA:cDNA Synthesis and Amplification (New England Biolabs, Cat #E6421). Further sample prep was performed with PacBio SMRTbell Express Template Prep Kit 2.0 (PacBio, PN 100-938-900) and Barcoded Overhang Adapter Kits A&B (Pacific Biosciences, Cat #PN 100-628-400 and #PN 100-628-500), with 1 µg of amplified cDNA per sample as input. The final library contained one barcoded cDNA sample per larval instar, for a total of five samples. The barcoded library was sequenced using the PacBio Sequel IIe platform. Consensus sequences for subreads were generated using ccs. All further subread processing was performed with the IsoSeq 3 pipeline (v4.0.0, https://github.com/PacificBiosciences/IsoSeq). Lima v2.7.1 was used to filter reads and remove primers and barcodes for each sample separately. Due to the low quantity of reads passing initial quality control, reads of different life stages were pooled for further analysis (**Supplementary Tables S2, S4**). PolyA tails and artificial concatemers were removed using the IsoSeq refine-function, after which reads were clustered using IsoSeq cluster. Clustered reads were mapped to the *A. luna* genome (GenBank accession number GCA_039707435.1) using pbmm2 v1.13.1, a SMRT C++ wrapper for minimap2 (Li 2018). Mapped reads were clustered into unique isoforms using IsoSeq collapse and subsequently classified and filtered using pigeon v1.2.0.

### Sericin Identification

A sericin dataset was generated from previously identified sericins from *A. selene* (Dong et al. 2015), *A. assamensis* (Dong et al. 2015), *A. pernyi* (Dong et al. 2015), *A. yamamai* (Žurovec et al. 2016), *B. mori* (Dong et al. 2019; Guo et al. 2022; Takasu et al. 2007), *H. cecropia* (Rouhová et al. 2024), *R. newara* (Dong et al. 2015), and *S. ricini* (*S. cynthia ricini*) (Dong et al. 2015; Tsubota et al. 2016). Each of these sequences were downloaded from the NCBI nucleotide database (Sayers et al. 2024), or from the original article when unavailable there. To increase the reliability of this sericin dataset, each putative sericin was compared to the NCBI nr database and sequences exhibiting high similarity with non-sericin sequences, such as *titin*, were excluded. Whenever a genome was available, duplicates were removed by only retaining the longest transcript matching to a particular genomic region. Using the final sericin dataset (**Supplementary Table S1**), sericins were discovered in the *A. luna* genome (GCA_039707435.1) using the command-line version of the Basic Local Alignment Search Tool (BLAST v2.14.1) (Altschul et al. 1990; Camacho et al. 2009) using both blastn and blastp with an Expect value threshold of 1e -50. Additionally, nucleotide sequences for saturniid sericins were aligned using muscle v5.1 (Edgar 2022) and matching patterns in the *A. luna* genome were identified with nhmmer (hmmer v3.4) (Wheeler & Eddy 2013). For each hit that contained a high number of serine-containing repeats, the surrounding 5,000 base pairs were extracted and imported into Geneious v 11.1.5 (https://www.geneious.com). Extracted genomic regions were manually annotated using the BRAKER3 automatic genomic annotation (Gabriel et al. 2024; Markee et al. 2024), the long-read silk gland specific transcriptome (see “Long-read Silk Gland Transcriptome”), and whole-body short-read RNA sequencing (see “Short-read RNA Sequencing and Read Alignment”) to identify the number of exons and intron-exon boundaries. Exons were deemed alternative exons if they were not present in all long read-transcripts or if short reads bridging the two neighboring exons were found. Subsequently, genomic regions around each sericin were scanned for recent duplications. The presence of a signal peptide was confirmed using the web service of SignalP 6.0 server (Teufel et al. 2022). Identification was confirmed with an online BLAST search to the NCBI nr database (Camacho et al. 2009; Sayers et al. 2024).

### Chromosome Synteny Maps

Genomic locations of sericins in the genomes of *A. luna*, *A. yamamai, B. mori,* and *S. ricini* (NCBI accession numbers, respectively: GCA_039707435.1; GCA_036509395.1; GCF_030269925.1; GCA_014132275.2) were compared using ChromSyn (Edwards et al. 2022). A set of 5286 single copy highly conserved orthologues sequences were identified using BUSCO v5.3.0 (Simão et al. 2015) with a lepidoptera-specific dataset. Telomere locations were identified and contig lengths were extracted for each genome with telociraptor v0.11.0 (https://github.com/slimsuite/telociraptor). Synteny maps were generated with ChromSyn in R v4.4 (R Core Team 2010).

### Gene Phylogenetic Network

Sericin sequences for *B. mori* and representative saturniid taxa (**Supplementary Table S1**, see “Sericin Identification”) were aligned using muscle v5.1 (Edgar 2022) using default settings. Pairwise distance matrices for each alignment were generated using P Distance (Hamming 1950). Gene phylogenetic networks were generated and visualized with SplitsTree App 6.0.0 (Huson & Bryant 2024), using the Neighbor Net method (Bryant & Huson 2023) to obtain splits.

### Repeat Sequence Motifs

Sericin sequences were imported in Geneious v 11.1.5 (https://www.geneious.com). Repeats were manually extracted and aligned using muscle v3.8.425. The resulting alignment was used to generate a sequence logo in Geneious, showing the consensus sequence as well as the relative amino acid frequency and information content (in bits) at every position (Schneider & Stephens 1990).

## Data Availability Statement

The data underlying this article are available as a bioproject in the GenBank Nucleotide Database at https://www.ncbi.nlm.nih.gov/genbank/, and can be accessed using the following bioproject accession number: PRJNA1072661. The protein alignment used for the “Gene Phylogenetic Network” and the genome feature file used for “Short-read RNA Sequencing and Read Alignment” are available on figshare at https://figshare.com/, and can be accessed at the following DOI: 10.6084/m9.figshare.29847707.

## Acknowledgements

We thank Paul Masonick and Bryce Shirk for providing valuable manuscript feedback. Gisella DePiazza, Nolan Ferguson, Ava Johnson, and Olivia Van Der Vlugt helped rear caterpillars. We acknowledge the use of HPC resources of UFIT Research Computing for all bioinformatic analyses.

