## Supplementary data for "Evolution of highly repetitive silk genes in the Luna moth, *Actias luna*"

**Table S1. Included sericin sequences.**

| Species | Name | Accession number | Reference |
|---|---|---|---|
| *Actias selene* | *Unigene3219* | GBZL01003169.1 | Dong et al. (2015) |
| *Actias selene* | *Unigene3639* | GBZL01003581.1 | Dong et al. (2015) |
| *Actias luna* | *Sericin 1* | PX095427 | Current study |
| *Actias luna* | *Sericin A* | PX095434 | Current study |
| *Actias luna* | *Sericin B* | PX095433 | Current study |
| *Actias luna* | *Sericin C* | PX095432 | Current study |
| *Actias luna* | *Sericin D* | PX095431 | Current study |
| *Actias luna* | *Sericin E* | PX095430 | Current study |
| *Actias luna* | *Sericin F* | PX095429 | Current study |
| *Actias luna* | *Sericin G* | PX095428 | Current study |
| *Antheraea assamensis* | *Unigene1892* | GBZC01001844.1 | Dong et al. (2015) |
| *Antheraea assamensis* | *Unigene11074* | GBZC01010713.1 | Dong et al. (2015) |
| *Antheraea pernyi* | *Unigene3413* | GBZF01003354.1 | Dong et al. (2015) |
| *Antheraea pernyi* | *Unigene13275* | GBZF01012995.1 | Dong et al. (2015) |
| *Antheraea pernyi* | *Unigene25271* | GBZF01024717.1 | Dong et al. (2015) |
| *Antheraea pernyi* | *Unigene3233* | GBZF01003179.1 | Dong et al. (2015) |
| *Antheraea yamamai* | *Sericin 1 (Src1)* | BBA53792.1 | Zurovec et al. (2016) |
| *Antheraea yamamai* | *Sericin 2 (Src2)* | BBA53793.1 | Zurovec et al. (2016) |
| *Antheraea yamamai* | *Sericin 3 (Src3)* | BBA53794.1 | Zurovec et al. (2016) |
| *Antheraea yamamai* | *Sericin 4 (Src4)* | BBA53795.1 | Zurovec et al. (2016) |
| *Antheraea yamamai* | *Sericin 5 (Src5)* | BBA53796.1 | Zurovec et al. (2016) |
| *Bombyx mori* | *Sericin 1 (Ser1)* | XP_037869538.1 | NA |
| *Bombyx mori* | *Sericin 2 (Ser2)* | NP_001166287.1 | NA |
| *Bombyx mori* | *Sericin 3 (Ser3)* | NP_001108116.1 | Takasu et al. (2007) |
| *Bombyx mori* | *Sericin 4 (Ser4)* | UTH80338.1 | Dong et al. (2019) |
| *Bombyx mori* | *Sericin 5 (Ser5)* | WFQ96466.1 | (Guo et al. 2022) |
| *Hyalophora cecropia* | *Sericin 1 (Src1)* | WWE94420.1 | Rouhova et al. (2024) |
| *Hyalophora cecropia* | *Sericin 2 (Src2)* | WWE94421.1 | Rouhova et al. (2024) |
| *Rhodinia newara* | *Unigene3473* | N/A | Dong et al. (2015) |
| *Rhodinia newara* | *Unigene12767* | GBZE01012589.1 | Dong et al. (2015) |
| *Samia ricini* | *Serine-rich protein 1* | LC001866.1 | Tsubota et al. (2016) |
| *Samia ricini* | *Serine-rich protein 4* | LC001869.1 | Tsubota et al. (2016) |
| *Samia ricini* | *Serine-rich protein 5* | LC001870.1 | Tsubota et al. (2016) |
| *Samia ricini* | *Unigene3618* | GBZD01003513.1 | Dong et al. (2015) |
| *Samia ricini* | *Unigene1738* | GBZD01001705.1 | Dong et al. (2015) |

**Table S2**. ***Actias luna* IsoSeq filtering statistics.** ZMW: zero-mode waveforms. flnc: full-length non-chimeric reads. L1-5: first – fifth instar caterpillars. The percentage of retained reads was calculated by dividing the number of retained reads by the input ZMWs.

| | L1 | L2 | L3 | L4 | L5 | Unbarcoded | Combined |
|---|---|---|---|---|---|---|---|
| Input ZMWs | 398483 | 403604 | 1214688 | 1118299 | 1140205 | 2944823 | 7220102 |
| ZMWs retained after lima filtering | 366290 | 363618 | 1046311 | 940923 | 958694 | 219540 | 3895376 |
| flnc retained after isoseq refine | 365192 | 362262 | 1021790 | 930422 | 950992 | 210606 | 3841264 |
| Reads retained after pigeon | 232628 | 226029 | 196036 | 257761 | 390330 | 19725 | 2299073 |
| Reads retained after pigeon (%) | 58.38 | 56.00 | 16.14 | 23.05 | 34.23 | 0.67 | 31.84 |

**Table S3**. ***Actias luna* short read RNAseq Metadata, adapted from Markee et al. (2024) .** “pre-pupa” is represented as fifth instar throughout the paper.

| **Sample ID** | **Sequence name** | **SRA experiment** | **SRA run** | **Life stage** | **Tissue type** |
|---|---|---|---|---|---|
| luna_01_head | 2017RNALibPool01-1 | SRX24005177 | SRR28400506 | 4th instar | head |
| luna_06_thorax | 2017RNALibPool01-10 | SRX24005178 | SRR28400505 | 4th instar | thorax |
| luna_06_abdomen* | 2017RNALibPool01-11 | SRX24005189 | SRR28400494 | 4th instar | abdomen |
| luna_01_thorax | 2017RNALibPool01-2 | SRX24005200 | SRR28400483 | 4th instar | thorax |
| luna_01_abdomen* | 2017RNALibPool01-3 | SRX24005209 | SRR28400474 | 4th instar | abdomen |
| luna_02_head | 2017RNALibPool01-4 | SRX24005210 | SRR28400473 | 4th instar | head |
| luna_02_thorax | 2017RNALibPool01-5 | SRX24005211 | SRR28400472 | 4th instar | thorax |
| luna_03_wholebody | 2017RNALibPool01-6 | SRX24005212 | SRR28400471 | 6 eggs | whole body |
| luna_04_wholebody* | 2017RNALibPool01-7 | SRX24005213 | SRR28400470 | 1st instar | whole body |
| luna_05_wholebody* | 2017RNALibPool01-8 | SRX24005214 | SRR28400409 | 1st instar | whole body |
| luna_06_head | 2017RNALibPool01-9 | SRX24005179 | SRR28400504 | 4th instar | head |
| luna_07_head | 2017RNALibPool02-1 | SRX24005180 | SRR28400503 | pre-pupa | head |
| luna_10_head | 2017RNALibPool02-10 | SRX24005181 | SRR28400502 | pupa | head |
| luna_10_thorax | 2017RNALibPool02-11 | SRX24005182 | SRR28400501 | pupa | thorax |
| luna_07_thorax | 2017RNALibPool02-2 | SRX24005183 | SRR28400500 | pre-pupa | thorax |
| luna_07_abdomen* | 2017RNALibPool02-3 | SRX24005184 | SRR28400499 | pre-pupa | abdomen |
| luna_08_head | 2017RNALibPool02-4 | SRX24005185 | SRR28400498 | pre-pupa | head |
| luna_08_thorax | 2017RNALibPool02-5 | SRX24005186 | SRR28400497 | pre-pupa | thorax |
| luna_08_abdomen* | 2017RNALibPool02-6 | SRX24005187 | SRR28400496 | pre-pupa | abdomen |
| luna_09_head | 2017RNALibPool02-7 | SRX24005188 | SRR28400495 | pre-pupa | head |
| luna_09_thorax | 2017RNALibPool02-8 | SRX24005190 | SRR28400493 | pre-pupa | thorax |
| luna_09_abdomen* | 2017RNALibPool02-9 | SRX24005191 | SRR28400492 | pre-pupa | abdomen |
| luna_11_head | 2017RNALibPool03-1 | SRX24005192 | SRR28400491 | pupa | head |
| luna_15_head | 2017RNALibPool03-10 | SRX24005193 | SRR28400490 | adult | head |
| luna_15_thorax | 2017RNALibPool03-11 | SRX24005194 | SRR28400489 | adult | thorax |
| luna_15_abdomen | 2017RNALibPool03-12 | SRX24005195 | SRR28400488 | adult | abdomen |
| luna_11_thorax | 2017RNALibPool03-2 | SRX24005196 | SRR28400487 | pupa | thorax |
| luna_11_abdomen | 2017RNALibPool03-3 | SRX24005197 | SRR28400486 | pupa | abdomen |
| luna_12_head | 2017RNALibPool03-4 | SRX24005198 | SRR28400485 | pupa | head |
| luna_12_thorax | 2017RNALibPool03-5 | SRX24005199 | SRR28400484 | pupa | thorax |
| luna_12_abdomen | 2017RNALibPool03-6 | SRX24005201 | SRR28400482 | pupa | abdomen |
| luna_14_head | 2017RNALibPool03-7 | SRX24005202 | SRR28400481 | adult | head |
| luna_14_thorax | 2017RNALibPool03-8 | SRX24005203 | SRR28400480 | adult | thorax |
| luna_14_abdomen | 2017RNALibPool03-9 | SRX24005204 | SRR28400479 | adult | abdomen |
| luna_16_wholebody* | 2017RNALibPool04-1 | SRX24005205 | SRR28400478 | 1st instar | whole body |
| luna_13_thorax | 2017RNALibPool05-8 | SRX24005206 | SRR28400477 | adult | thorax |
| luna_02_abdomen* | 2017RNALibPool07-1 | SRX24005207 | SRR28400476 | 4th instar | abdomen |
| luna_13_abdomen | 2017RNALibPool07-10 | SRX24005208 | SRR28400475 | Adult | abdomen |

** Indicates which samples were used in the differential expression analysis.*

**Table S4**. ***Actias luna* IsoSeq Metadata. "**1$^{st}$-5$^{th}$" instar represents unbarcoded reads. SRA = Sequence Read Archive

| **Sample ID** | **SRA experiment** | **Life stage** | **Tissue type** |
|---|---|---|---|
| * SAMN49719316 | SRX29471812 | 1$^{st}$ instar | Silk gland |
| * SAMN49719317 | SRX29471813 | 2$^{nd}$ instar | Silk gland |
| * SAMN49719318 | SRX29471814 | 3$^{rd}$ instar | Silk gland |
| * SAMN49719319 | SRX29471815 | 4$^{th}$ instar | Silk gland |
| * SAMN49719320 | SRX29471816 | 5$^{th}$ instar | Silk gland |
| SAMN49719321 | SRX29471817 | 1$^{st}$-5$^{th}$ instar | Silk gland |

** Indicates which samples were used to generate long read transcriptome.*

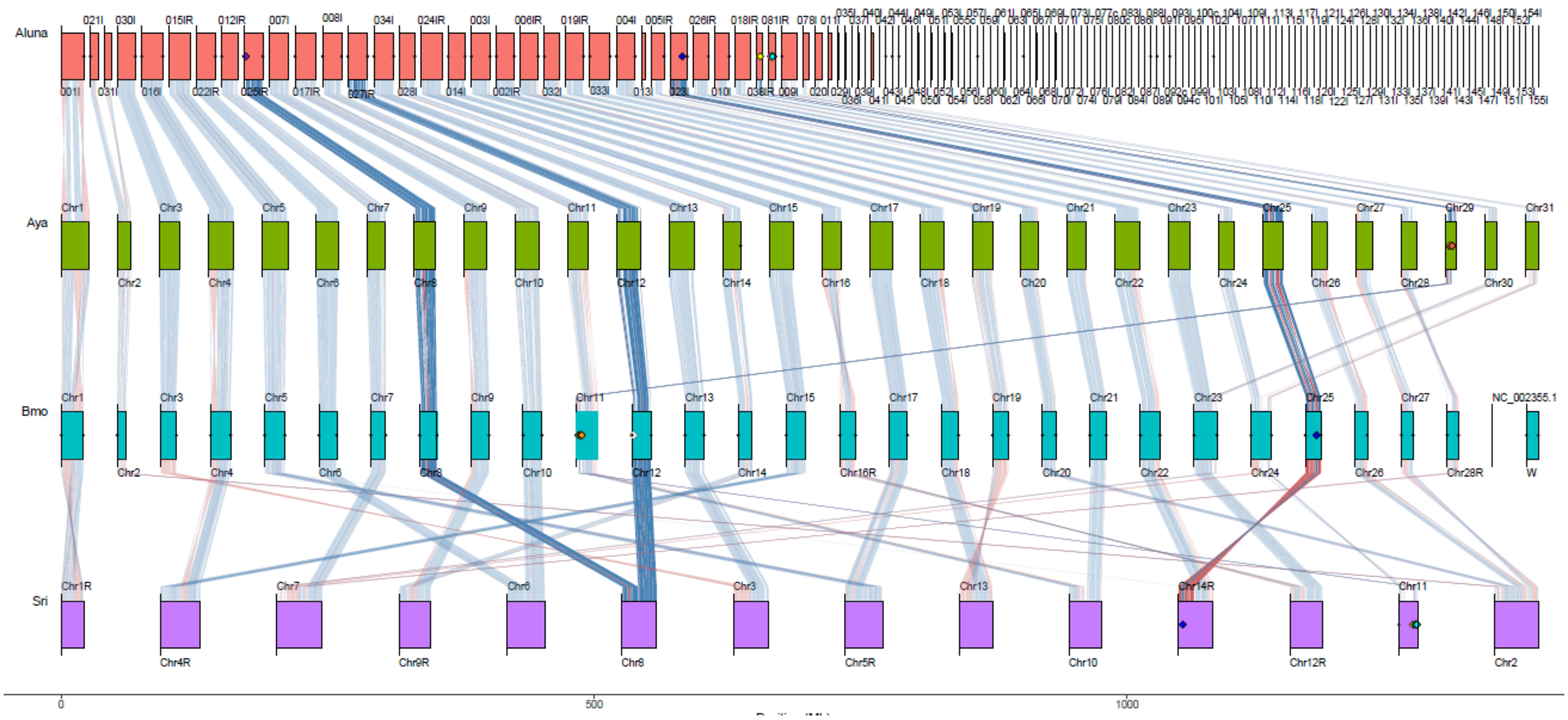
Aluna
Aya
Bmo
Sri
0
500
1000

**Figure S1. Chromosome synteny map**. ChromSyn BUSCO-derived synteny plots of the chromosomes for four different species of moths (*A. luna, A. yamamai, S. ricini, B. mori*). Small black filled circles at either end of a chromosome mark telomere predictions. Filled diamonds mark *FibH* and sericin locations (blue: *FibH*, yellow: *A. yamamai src1*, *A. luna ser1*, *S. ricini unigene2618*, *B. mori ser1;* red: *A. luna serE-G*, *A. yamamai src2-5*, *S. ricini srp4-5*; cyan: *A. luna serB-D*, *S. ricini srp1*, *S. ricini unigene1738*, *B. mori ser5*; purple: *A. luna serA*, white: *B. mori sericin P150*, orange: *B. mori ser2-4*). Contig names were retained from the respective NCBI assemblies but "ptg000" was trimmed from each of the *A. luna* contigs; an "R" denotes contigs reversed in orientation. Blue and red lines depict synteny blocks of BUSCO genes (blue, same strand; red, inversion), with darker lines connecting chromosomes or chromosome sections that contain sericins or FibH and fainter lines connecting the remaining chromosomes. Aya = *A. yamamai*, Bmo = *B. mori*, Sri = *S. ricini*

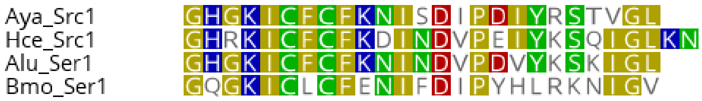


**Figure S2. Alignment of sericin 1 N-termini.** N-termini were extracted from sericin 1 proteins from *A. yamamai* (Aya), *H. cecropia* (Hce), *A. luna* (Alu) and *B. mori* (Bmo). Conserved residues were colored based on their polarity (blue: basic, red: acidic, green: polar, yellow: non-polar).

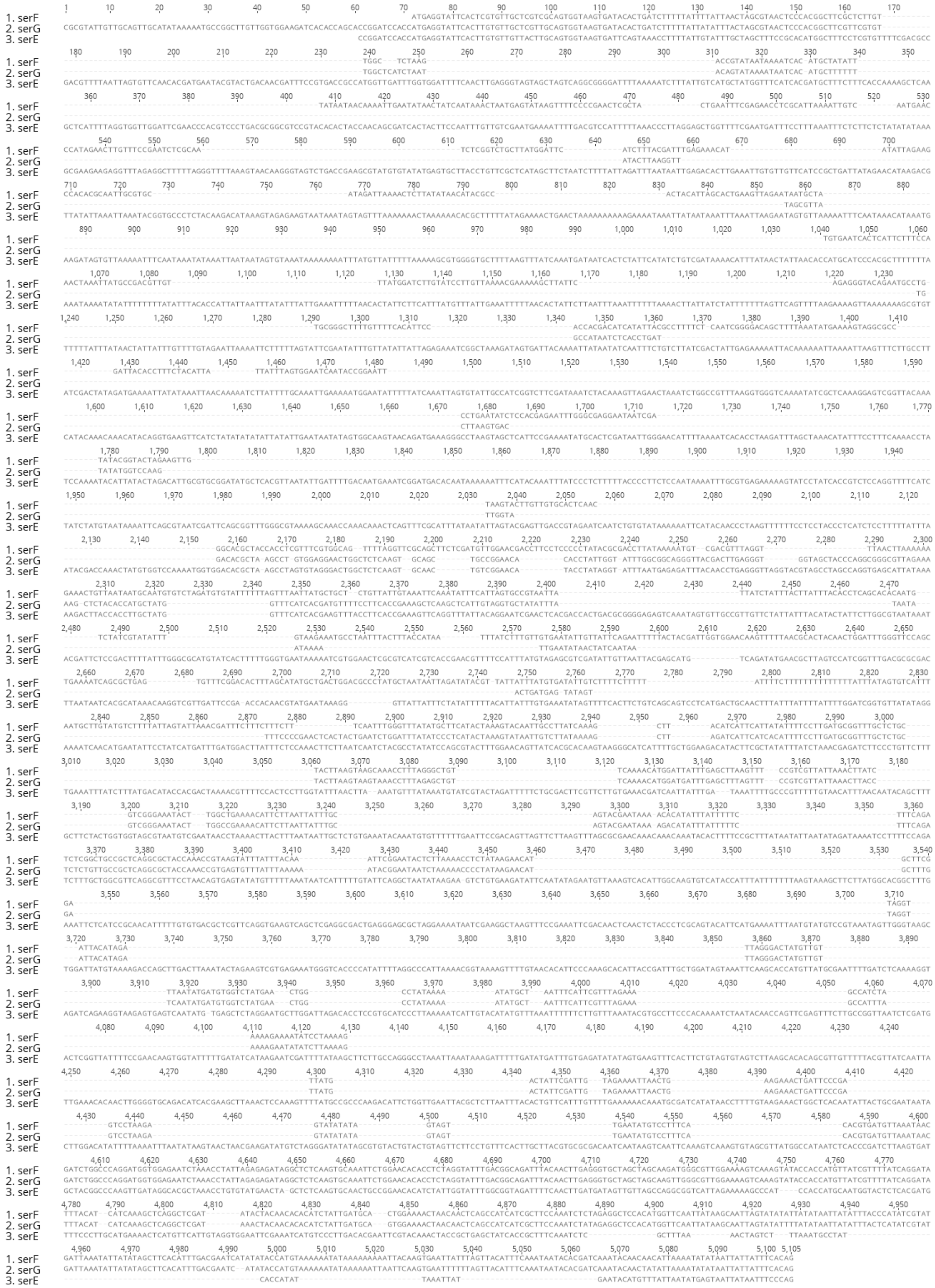


**Figure S3. Alignment of C-terminus of *A. luna* Group 3 sericins.** C-termini, including the first two exons and two introns, were extracted from *A. luna* Group 3 sericins (serE,F,G) and aligned using muscle.
